# A low-cost device for cryoanesthesia of neonatal rodents

**DOI:** 10.1101/2022.06.09.495437

**Authors:** Bradley B. Jamieson, Xavier Cano-Ferrer, George Konstantinou, Elisa de Launoit, Nicolas Renier, Albane Imbert, Johannes Kohl

**Affiliations:** State-dependent Neural Processing Laboratory, The Francis Crick Institute, 1 Midland Rd, London NW1 1AT, UK; Making STP, The Francis Crick Institute, 1 Midland Rd, London NW1 1AT, UK; Sorbonne Université, Paris Brain Institute - ICM, INSERM, CNRS, AP-HP, Hôpital de la Pitié Salpêtrière, Paris, France

**Keywords:** Cryoanesthesia, Rodents, Neonates, Peltier, Stereotaxic surgery, Thermoelectric device

## Abstract

Studying the development of neural circuits in rodent models requires surgical access to the neonatal brain. Since commercially available stereotaxic and anesthetic equipment is designed for use in adults, reliable targeting of brain structures in such young animals can be challenging. Hypothermic cooling (cryoanesthesia) has been used as a preferred anesthesia approach in neonates. This commonly involves submerging neonates in ice, an approach that is poorly controllable. We have developed an affordable, simple to construct device – CryoPup – that allows for fast and robust cryoanesthesia of rodent pups. CryoPup consists of a microcontroller controlling a Peltier element and a heat exchanger. It is capable of both cooling and heating, thereby also functioning as a heating pad during recovery. Importantly, it has been designed for size compatibility with common stereotaxic frames. We validate CryoPup in neonatal mice, demonstrating that it allows for rapid, reliable and safe cryoanesthesia and subsequent recovery. This open-source device will facilitate future studies into the development of neural circuits in the postnatal brain.

**Specifications table:** 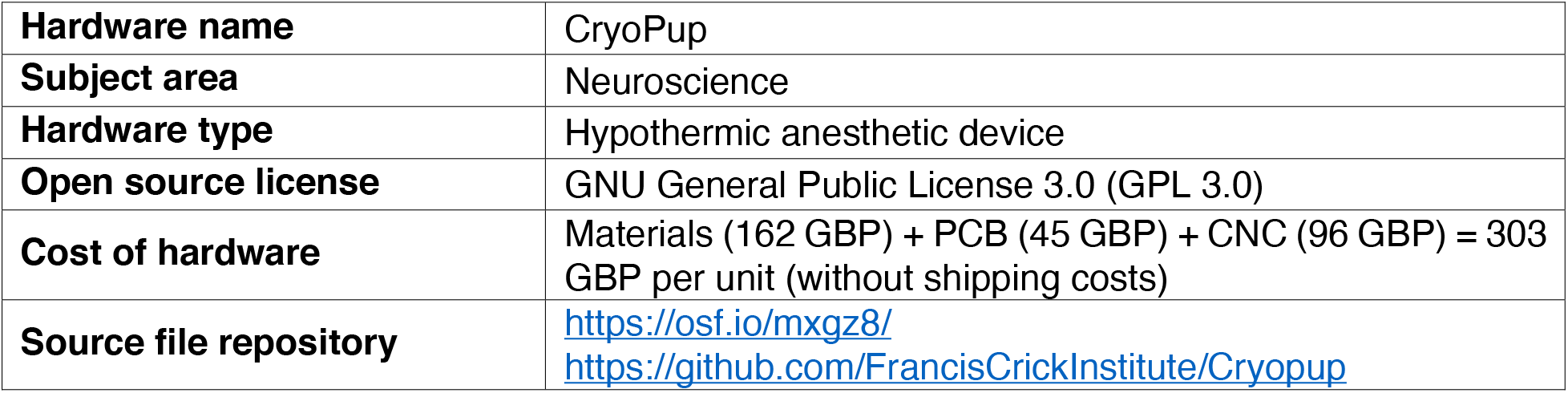

## 1. Hardware in context

Systems neuroscience research heavily relies on visualizing and manipulating defined neural circuits in the brain. In rodent models, this is commonly achieved via precise microinjection of chemicals, implantation of cannulas, electrodes and optic fibers, or delivery of viral vectors into the brain using stereotaxic surgery [1,2]. For this purpose, animals are secured into a stereotaxic frame while anesthetics are delivered via intraperitoneal injection or inhalation through a mouthpiece [3].

While this procedure is routinely used in adults, it is suboptimal for neonates, which are increasingly used to study how neural circuits develop postnatally [4–8]: inhalation anesthetics can negatively and lastingly affect the brain of neonatal rodents, in particular areas associated with memory [9–12]. Instead, cryoanesthesia, i.e. anesthesia by deep hypothermia, has been commonly used in such young animals. This is typically carried out by submerging neonates in crushed ice or using simple cooling pads [7,8,13–15]. These approaches, however, do not provide accurate or consistent temperature control, and questions about the safety and appropriateness of hypothermic anesthesia for neonates have thus been raised [16]. A system capable of adjustable temperature control for hypothermic surgeries in adult rodents was recently published, but it is incompatible with stereotaxic surgeries due to its dimensions [17].

We have designed a device (CryoPup) to overcome these limitations. Based on a bidirectionally controlled Peltier element, CryoPup is capable of rapid cooling and heating (temperature range 0ºC to +42ºC) at ambient temperature (21ºC), thus also acting as a heating pad (see section 7 for safe temperatures for cryoanesthesia and recovery). Due to its compact size, it is fully compatible with modern stereotaxic equipment. We have also designed a 3D printed head holder which facilitates the positioning of neonates during stereotaxic surgeries, thus increasing targeting success. Finally, we show that CryoPup allows for rapid, reliable and safe cryoanesthesia in neonatal mice.

This device thus represents a cost-effective and versatile solution for cryoanesthesia in neonatal rodents.

## 2. Hardware description

CryoPup combines a thermoelectric module (Peltier element) which rapidly cools to temperatures suitable for cryoanesthesia of neonatal rodents, with 3D printed components to form a surgery bed. The device consists of a thermoelectric element which pumps heat from a low to a high temperature side and a CPU water cooling system which removes heat from the high temperature side. Peltier-based cooling provides accurate temperature control for cryoanesthesia in a closed loop with a thermocouple confined between the bed top plate and the Peltier element. Temperature is set via an intuitive control panel and the device is compatible with standard stereotaxic frames without further modifications. A 3D printed head holder ensures cranial stability during stereotaxic surgeries. Importantly, CryoPup can seamlessly switch from cooling to heating, and thus acts as a heating pad during recovery, or during the use of other forms of anesthesia in adults. This device can in principle also be equipped with a temperature probe to monitor external or internal (e.g. rectal) temperature.

### Main components

The device is composed of a main printed circuit board (PCB), the heat removing system and the bed which encloses a thermoelectric element (Fig. 1a-b).

**Fig. 1.**
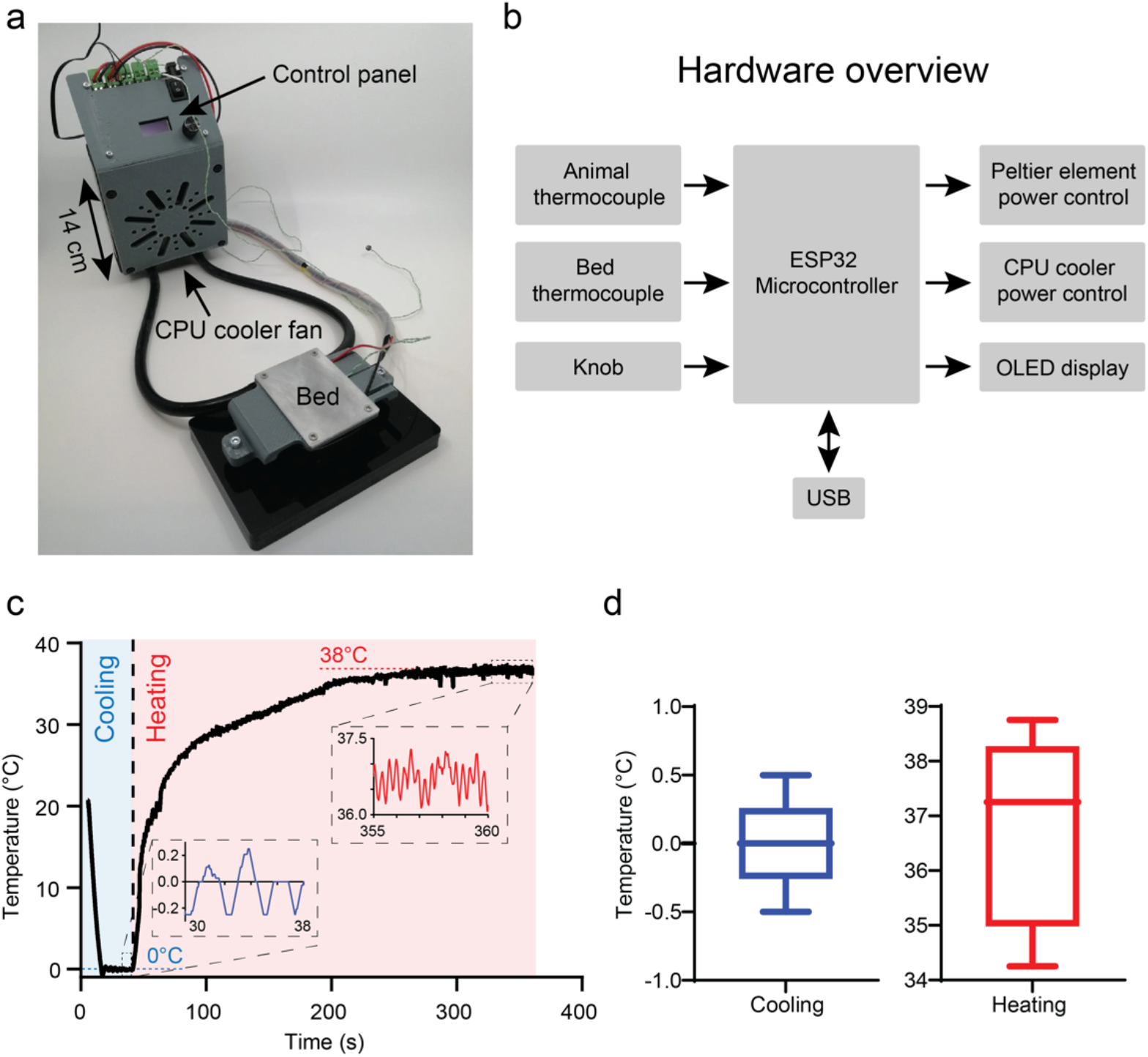
CryoPup device (a) and schematic control diagram (b). Device performance in a cooling (target temp. 0°C) and heating (target temp. 38°C) cycle (c). Temperature variation around the setpoints (c). Data (850 data points each) are from the trial shown in (c). Box-and-whisker plots show median, interquartile range and range. Note that temperatures of 0°C and 38°C are not safe for use in live animals.

### Electronics

The PCB hosts a microcontroller (Espressif ESP32) which controls the Peltier element and heat exchanger (CPU water cooling loop). Two Maxim integrated thermocouple amplifiers and digital converters (MAX31855KASA+T) acquire data from two Type-K temperature sensors: one in the bed and one external sensor which can be used to monitor experimental animals’ temperature. Two power MOSFETs (IRFH8318TRPBF) control the power to the All-In-One liquid cooler (Corsair Hydro H60) and the thermoelectric element (TEC1-12709). A rotary encoder is used to set the desired bed temperature. The desired temperature as well as the two thermocouple temperatures are continuously displayed on a 0.96-inch OLED display (Fig. 5b).

### Firmware

CryoPup is capable of both cooling and heating. Rapid cooling is achieved by the thermoelectric element pumping heat over a short distance and by the highly effective heat removal by the closed loop liquid cooler (Fig. 1c). Bed temperature is adjusted combining two bang-bang control algorithms. When cooling (i.e. desired bed temperature is below a predefined ambient temperature) the microcontroller modulates power to the Peltier element using a bang-bang control while the heat removing system is constantly ON. In contrast, if a higher-than-ambient desired temperature is selected, the heat removal system shuts down and a bang-bang control with a small pulse width modulation (PWM) duty-cycle value is initiated. With the thermoelectric element operating in its inefficient regime, heat gradually accumulates on the bed. The oscillations (ringing) around the set temperature are bigger when warming (± 1.5°C) than cooling (± 0.5°C) – thus precise temperature control is ensured during cryoanesthesia (Fig. 1d).

## 3. Design files

The design files provide all necessary content to manufacture, assemble and use CryoPup.

**Table 1.** List of design files

| Design filename | File type | Open source license | Location of the file |
| --- | --- | --- | --- |
| Control board | Easy EDA/ Gerber/csv | GPL-3.0 License | OSF |
| Peltier holder | Autodesk Inventor/STL/Step | GPL-3.0 License | OSF |
| Bed plate | Autodesk Inventor/STL/Step | GPL-3.0 License | OSF |
| Electronics holder | Autodesk Inventor/STL/Step | GPL-3.0 License | OSF |
| Base | Autodesk Inventor/DXF/STL/Step | GPL-3.0 License | OSF |
| Base spacer | Autodesk Inventor/DXF/STL/Step | GPL-3.0 License | OSF |
| Head holder | STL/Step | GPL-3.0 License | OSF |

Control board: Design and manufacturing files for the PCB as well as Pick and Place (PnP) and Bill of Materials (BOM) files for turn-key PCB manufacturing and assembly.

Peltier holder: Design and manufacturing files for 3D printing the bed frame.

Bed plate: Design and manufacturing files needed for machining the aluminium bed plate.

Electronics holder: Design and manufacturing files for 3D printing the control panel frame which is attached to the heat exchanger and holds the PCB.

Base: Design and manufacturing files for 3D printing or laser cutting the bed base.

Base spacer: Design and manufacturing files for 3D printing or laser cutting the base spacer for the bed.

Head holder: Design and manufacturing files for 3D printing the neonate stereotaxic head holder.

## 4. Bill of materials

The components required to build CryoPup, and their costs, are listed in Table 2.

**Table 2.**
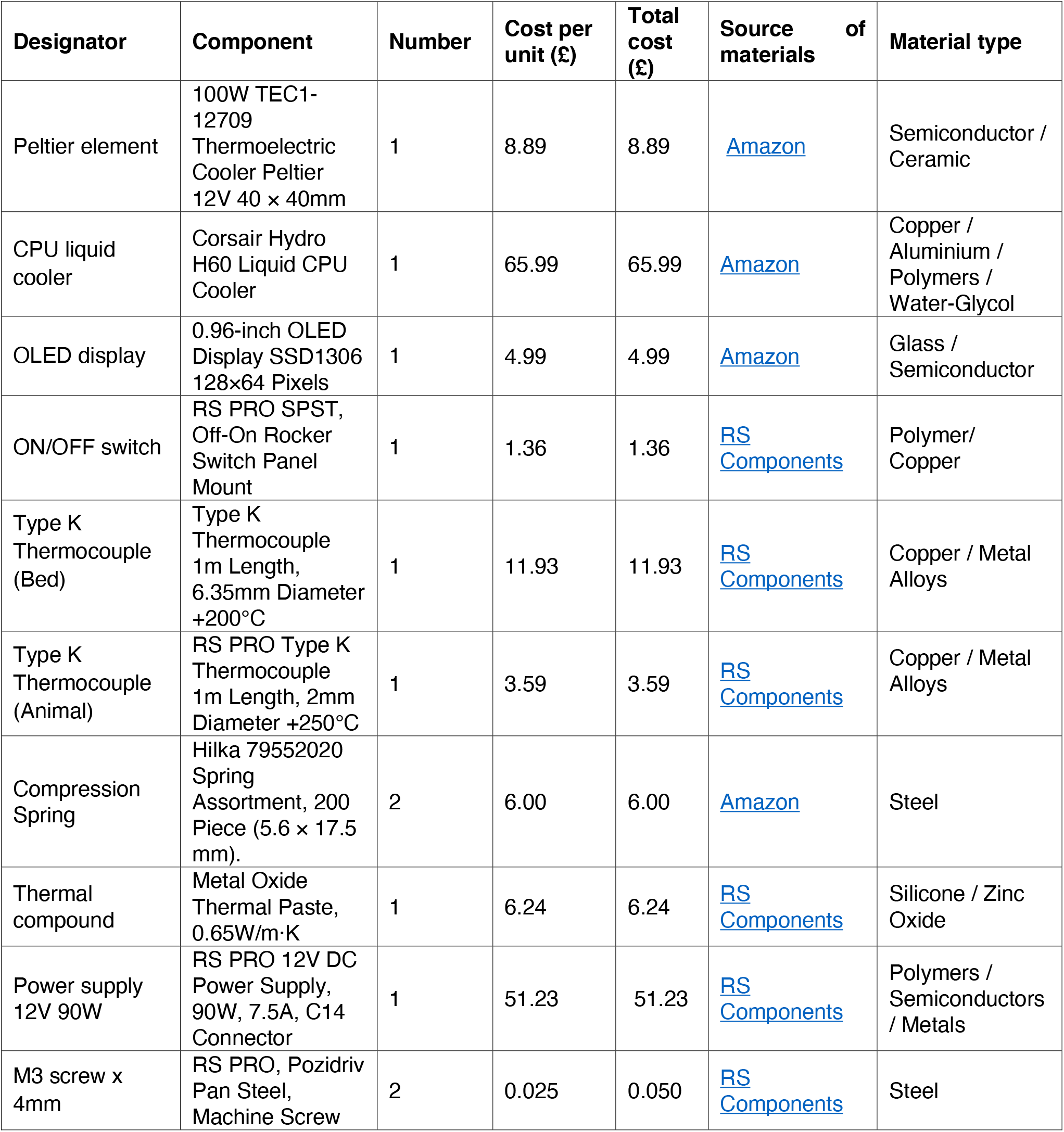
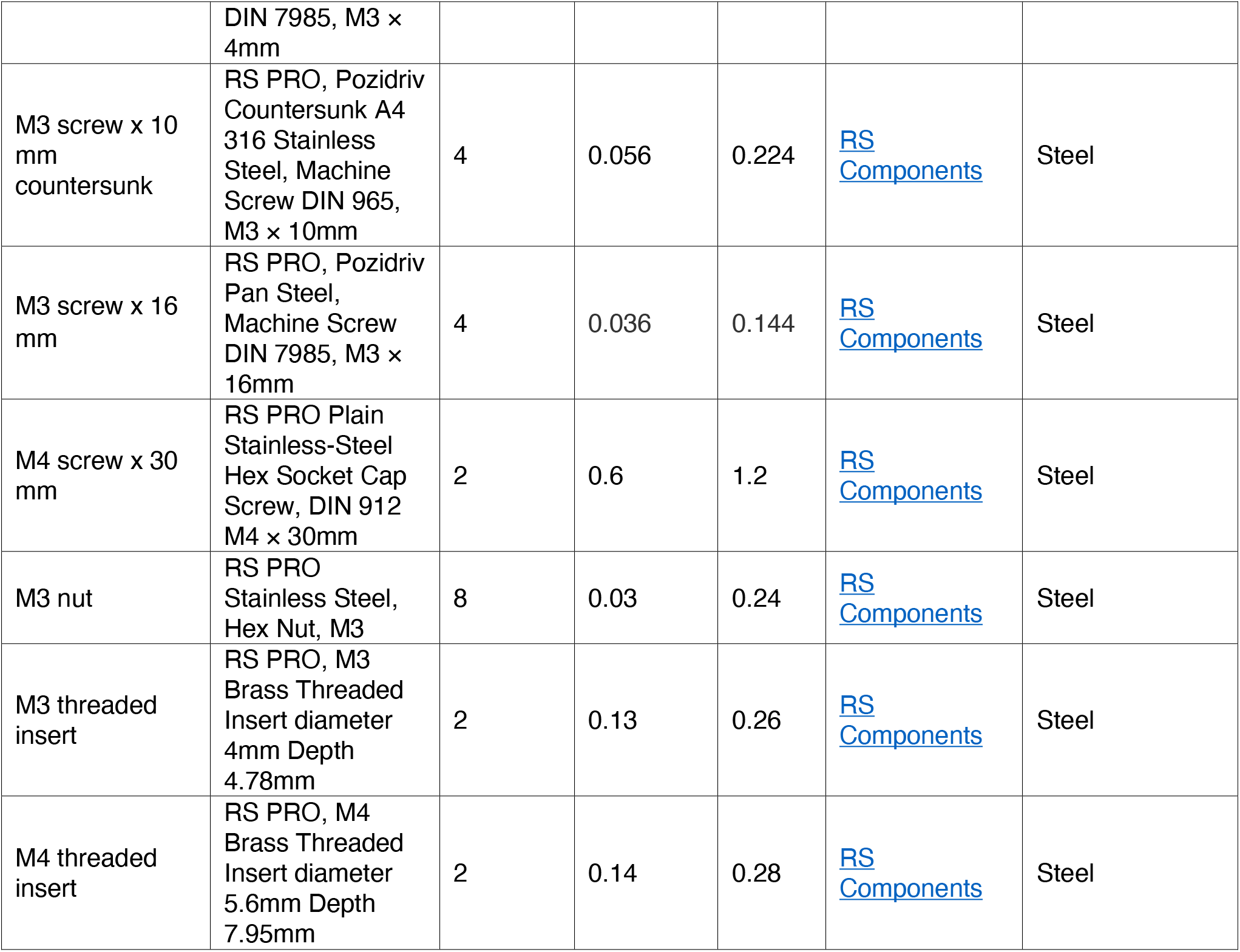
Components required for CryoPup

## 5. Build instructions

This section describes the build instructions for the PCB, bed base and electronics holder.

### 5.1. Required tools

1. FDM 3D printer (a Zortrax M200 with 200 × 200 × 180 mm building volume was used here)
2. Phillips screwdriver
3. Hex key set
4. Pliers
5. Side cutters / Cable strippers
6. Soldering iron and solder

### 5.2. Manufacturing the custom-designed PCB

The following manufacturing and assembly files for the PCB are available on the OSF repository (<u>link)</u>: Gerber files which contain the computer-aided manufacturing files, the bill of materials (BOM) that contains the part number associated with each component and the pick and place file (PnP) which specifies the coordinates on where each component must be placed by the pick and place machine. Following the simple steps outlined below, the board can be reproduced by the manufacturer (JLCPCB) without the need for ordering or soldering components.

1. Download the files from the OSF repository: <u>link</u>.
2. Create customer account on <u>JLCPCB</u> and sign in.
3. Go to <u>JLCPCB instant quote</u> and add Gerber file.
4. Select default settings and preferred PCB color.
5. Under PCB settings, select SMT assembly and indicate quantity (2 or 5).
6. Click next and add the BOM and PnP files.
7. Click next and review if any parts are unavailable. If yes, order from a different supplier (e.g. Mouser, Farnell, Digikey …).
8. Click next to see the position of each part on the PCB; save to cart and complete payment.

A more detailed document describing the procedure with images can be found on the repository manufacturing files folder (<u>link</u>).

### 5.3. Procuring the remaining components

A comprehensive Bill of Materials can be found in section 4.

### 5.4. Manufacturing of bed base and electronics holder

The four components to be 3D printed can be found in the <u>manufacturing files/mechanical parts</u> folder. The base and the base spacer parts can also be 3D printed but were laser cut from 10 mm acrylic sheets here. M4 and M3 threaded inserts are installed in the base and the Peltier holder parts (Fig. 2a).

**Fig. 2.**
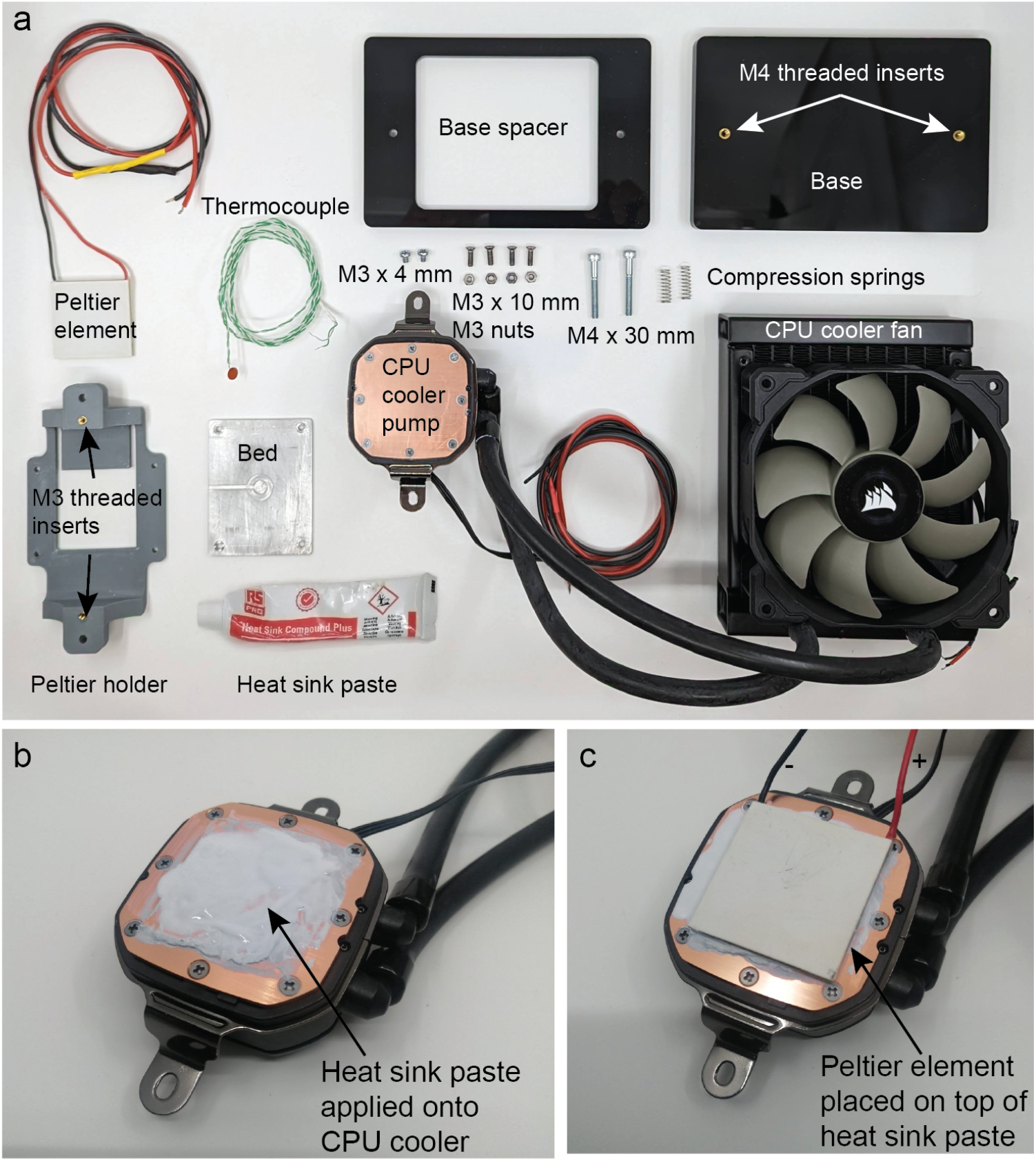
Required parts for the bed build (a). Application of thermal compound to top of cooler block (b). Peltier element placed on top of thermal compound with cold side facing up (location of red and black wires used as reference) (c).

### 5.5. Manufacturing the bed

The bed is manufactured from aluminium to ensure good temperature surface homogeneity. This was outsourced to a third-party manufacturer (<u>Hubs</u>), using standard settings (aluminium 6082, no additional treatments). The step files in the <u>Manufacturing files/Mechanical parts</u> folder can be directly uploaded to the instant quote webpage.

### 5.6. Bed assembly

After manufacturing all bed components (Fig. 2a), thermal compound is applied to the CPU cooler surface. Thermal compound improves the heat removal efficiency of the CPU cooler (Fig. 2b). The thermoelectric element is installed on top of the heat sink compound with the writing (hot side) facing the CPU cooler. By convention when the red cable of a thermoelectric element is connected to source positive and the black cable to negative, the ceramic plate on which the cables are soldered on is the hot side. Testing before installation is recommended.

The 3D printed Peltier holder is placed around the Peltier element (Fig. 3a) and attached to the CPU cooler using two 4 mm long M3 screws (Fig. 3b). The bed thermocouple is inserted into groove of the bed aluminium plate (Fig. 3c), the bed plate placed on top of the Peltier element enclosing the thermocouple and secured with 10 mm long countersunk M3 screws and nuts (Fig. 3d-e). The bed is attached to the base and base spacer using two M4 screws with a compression spring between the base and the Peltier holder (Fig. 3f-h). These springs allow for easy adjustment of bed height.

**Fig. 3.**
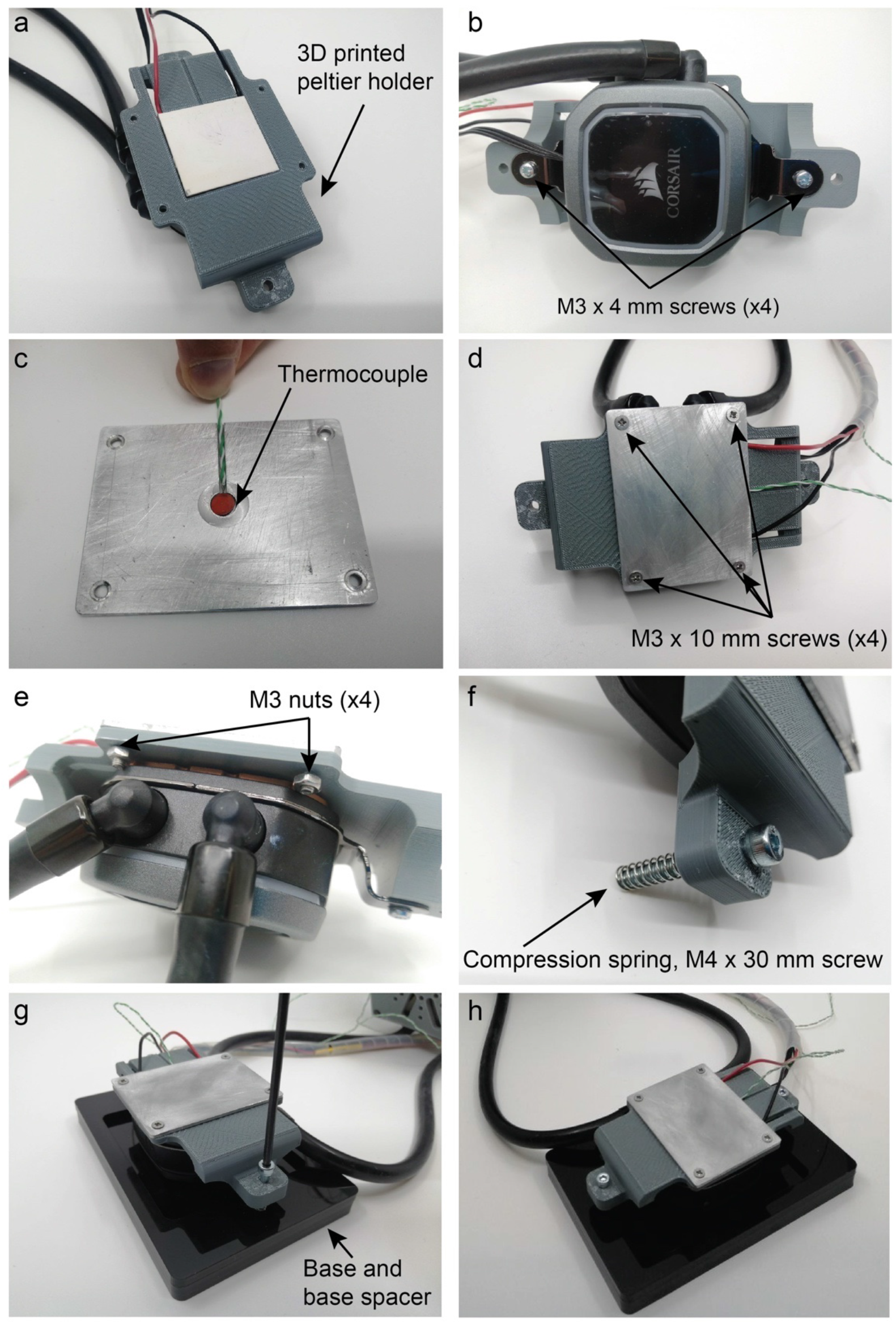
Installation of the thermoelectric module in the holder (a). Peltier holder attached to CPU cooling block (b). Thermocouple placed between Peltier element and bed aluminium plate (c). Aluminium plate attached to Peltier holder (d). Bed plate and Peltier holder (e). Compression springs provide counterforce to adjust height and to level the bed (f). Bed attached to laser cut base and base spacer (g). Finished assembly (h).

### 5.7. Electronics holder assembly

The OLED display and the ON/OFF switch are the final components to be soldered on the board. It is recommended to apply an isolating layer (electric or polyimide tape) to the back of the display before soldering it on the board (Fig. 4a).

**Fig. 4.**
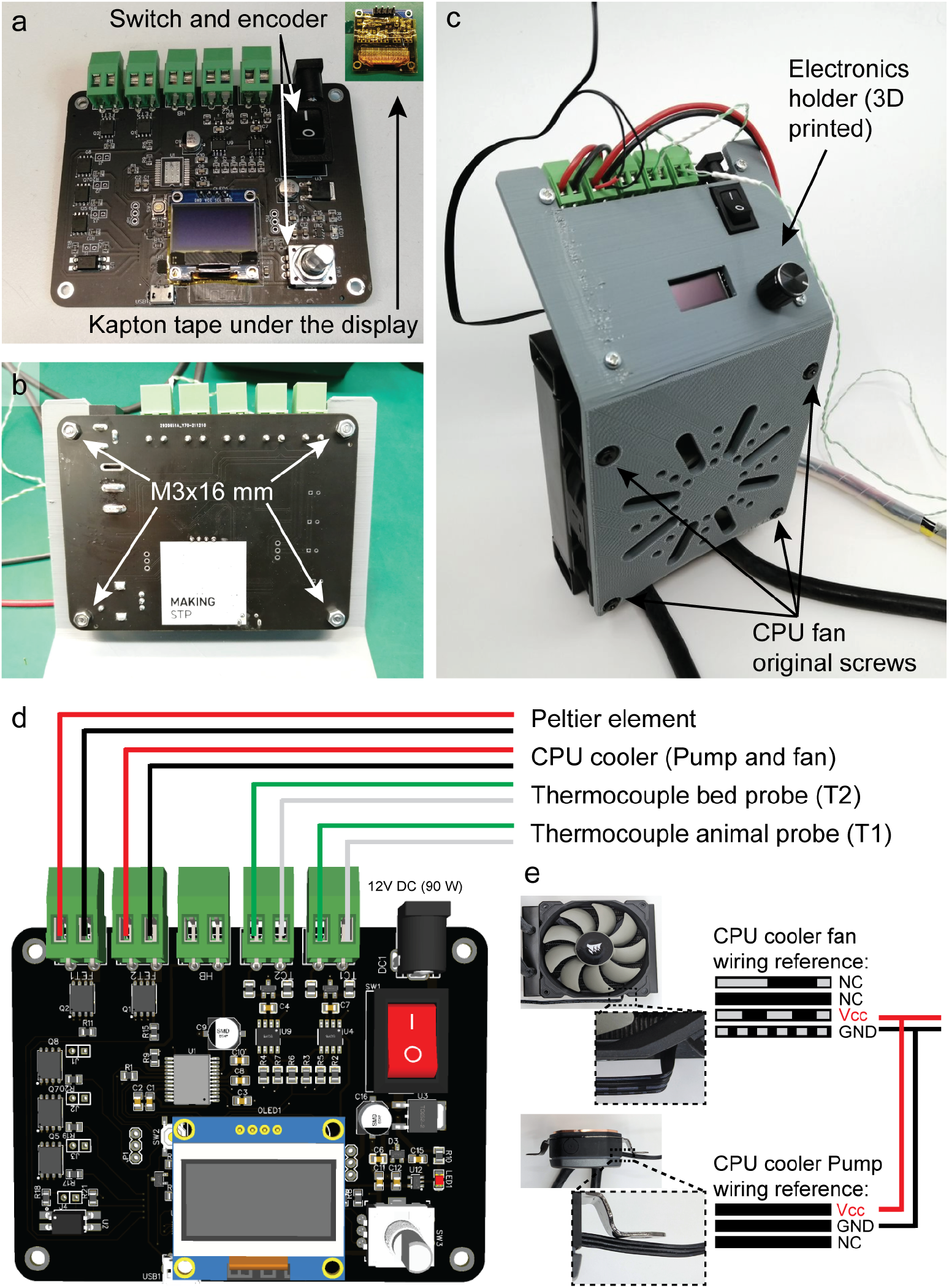
PCB with components and insulation between back of screen and microcontroller housing. PCB is fixed to the 3D printed electronics holder 2(1b4). Electronics holder attached to heat exchanger using the original fan screws (c). Electrical connections to/from PCB screw terminals (d). Heat exchanger fan 2a1n5d pump wire connections (e).

The firmware must be uploaded to the board before assembling it in the 3D printed electronics holder since the micro-USB port cannot be accessed afterwards. To upload the firmware on the board, follow the next steps (Windows):

1. Install the Arduino IDE on your computer (<u>link</u>)
2. If necessary, install the SiLabs CP2104 drivers (<u>link</u>)
3. Install the ESP32 package for the Arduino IDE (Version 1.0.6 recommended) (<u>link</u>)
4. Download the sketch from the repository (<u>link</u>)
5. Download libraries:
  i. ESP32_analog_write (<u>link</u>)
  ii. Encoder.h (<u>link</u>)
  iii. Adafruit_MAX31855.h (<u>link</u>)
  iv. Adafruit_GFX.h (<u>link</u>)
  v. Adafruit_SSD1306.h (<u>link</u>)
6. Connect the CryoPup board to the computer using a micro-USB cable and check “Device Manager” for the name of the COM port.
7. Select the board on Tools -> boards -> ESP32 Arduino -> Adafruit ESP32 Feather
8. Select the board on Tools -> ports -> the COM port listed in Device Manager.
9. Open the sketch *Cold_bed_2*.*ino* and upload it to the board (<u>link</u>)

After uploading the firmware, the PCB is attached to the 3D printed electronics holder using four 16 mm long M3 screws and nuts (Fig. 4b). The original fan screws are subsequently used to attach the electronics holder to the fan shroud (Fig. 4c). The frame also serves as a finger (and cable) guard. Finally, all electrical connections (for two thermocouples, CPU fan and pump, and Peltier element) are completed as specified in Fig. 4d. Since the CPU closed loop water cooler consists of two main elements (heat exchanger, i.e., fan and radiator, and pump/cooling block attached with the bed) which have different wiring connectors, they must be re-wired and connected to the same screw terminal as outlined in Fig. 4e.

### 5.8. Using the external thermocouple as a temperature probe (optional)

The external thermocouple can in principle be used as an internal (rectal) probe to monitor animal temperature in adults. For this purpose, the end of the external thermocouple probe can be covered with a 2cm piece of Tygon® E-3603 (3.2mm ID). This device is implantation biocompatible (USP class VI). However, we have not validated this functionality in animals.

## 6. Operation instructions

This section describes detailed steps for operating CryoPup.

1. Connect the 12 V power supply to the barrel jack (Fig. 5a, b).
2. Power up using the ON/OFF switch (Fig. 5b).
3. The display will show three parameters: the temperature from the external thermocouple (T1), the temperature from the bed (T2) and the desired temperature of the bed (DT). DT can be adjusted by the user at any time using the selection knob (default = 0°C).

**Fig. 5.**
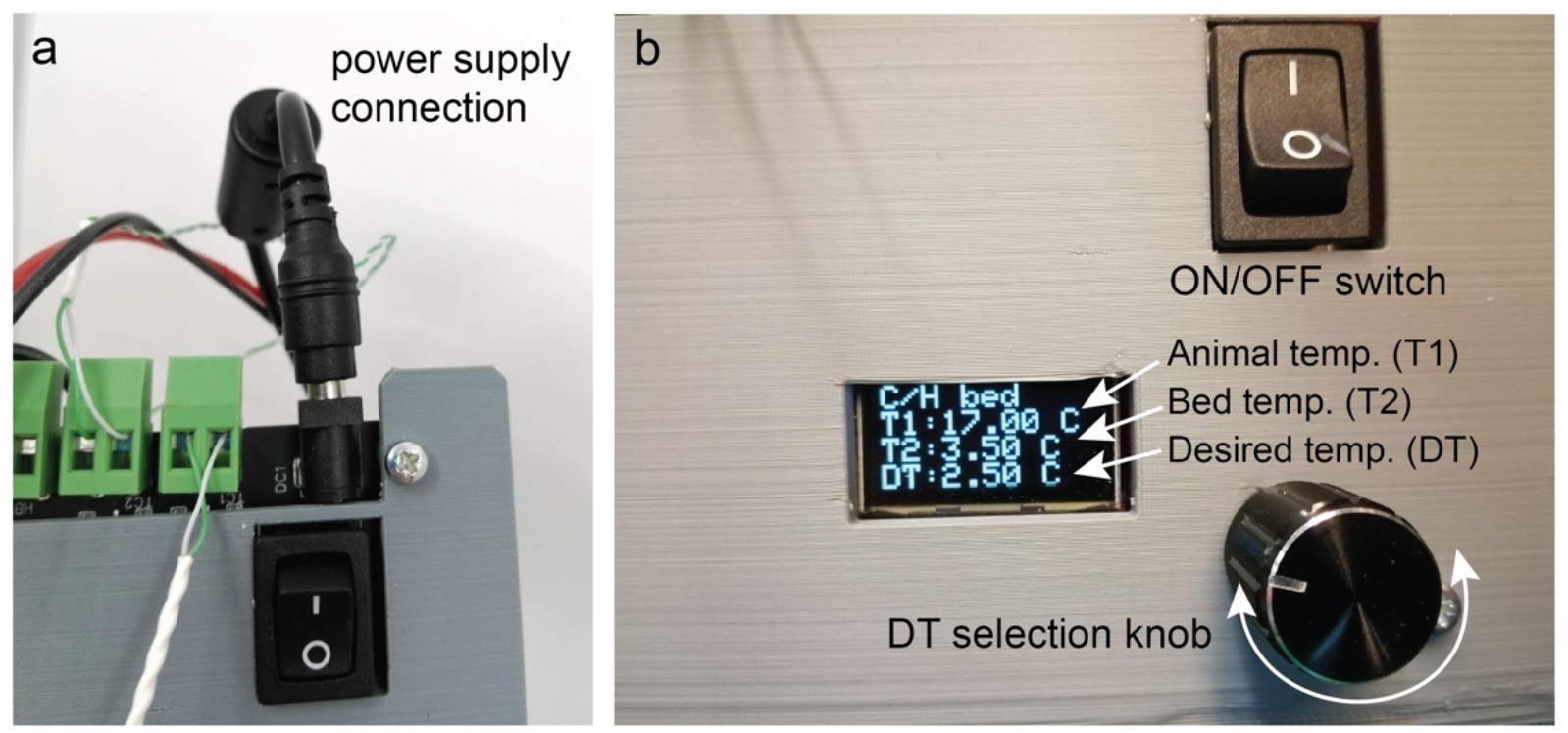
CryoPup is powered from a 12V DC 90W center positive power supply via a standard 2mm DC barrel jack connector (a). CryoPup control panel (b).

## 7. Validation and characterization

To test the functionality of CryoPup, we used it to anesthetise neonatal mice (postnatal day 2-3, n = 9) (Fig. 6a). For this purpose, the CryoPup device was slotted into a commonly used stereotactic frame (Kopf, Model 942). A 3D printed head holder (optional, see Table 1) was inserted into the frame’s mouse animal adaptor (Model 926, Traditional, Fig. 6a, b). CryoPup was set to a desired temperature (DT) of 2ºC to ensure a bed temperature (T2) of 4°C. A DT of 2ºC is necessary to achieve a bed temperature of 4ºC due to the temperature difference between the ambient temperature and the temperature measured by the thermocouple, which is directly in contact with the Peltier element. This 2ºC temperature difference is valid for lab environments which are temperature-controlled at 20-22ºC. We chose a target of 4°C since this temperature reliably induced cryoanesthesia. This is warmer than immersion of pups in ice (0ºC) or in water containing crushed ice (2-3ºC) [13], thus reducing the risk of adverse effects. Mouse pups were gently placed onto the device bed with surgical tape to ensure that the bed was in close contact with the pups’ chest/abdomen (Fig. 6a), and the blunt ends of standard ear bars (Kopf #1921) were used to gently fix the head in place (Fig. 6a, b). The pedal withdrawal reflex was tested every 30s with blunt forceps on alternating feet, and mice were assessed to have a level of anesthesia appropriate for surgery (surgical plane) when this reflex was absent from both feet. This was achieved in 123.8 ± 8.3 s (mean ± SEM; Fig. 6c, e).

**Fig. 6.**
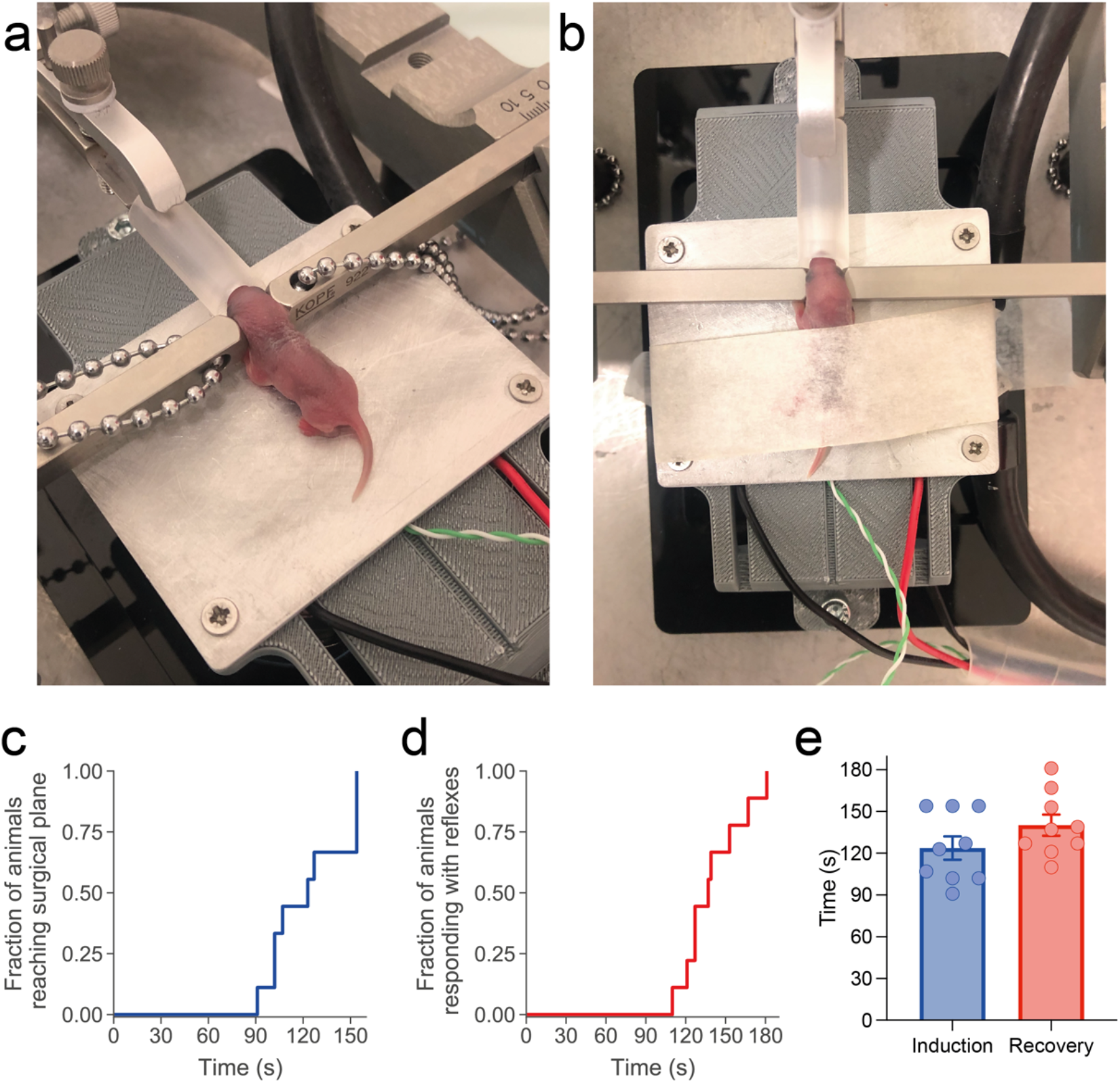
Example images of neonatal mice during cryoanesthesia on CryoPup (a, b). Survival curves showing the time of induction to reach surgical plane (c), and recovery rate following cessation of cryoanesthesia (d). Average induction and recovery latencies (e). Data presented as mean ± SEM.

Following induction, pups were maintained at 4°C for 15 minutes before DT was set to room temperature (21°C). The pedal withdrawal reflex was again monitored, and recovery was considered complete when this reflex had returned (140.2 ± 7.6 s, mean ± SEM; Fig. 6d, e). Animals were subsequently transferred to a standard heated recovery chamber to fully assess their mobility before returning them to the dam.

CryoPup is thus capable of (1) rapidly inducing cryoanesthesia, (2) stably maintaining this state for periods of time compatible with surgical procedures such as intracranial stereotaxic injections (which can be performed in ∼5min per coordinate in our experience), and (3) safely returning experimental subjects to wakefulness. Since both the control unit and the fan are potentially significant sources of electrical noise, appropriate grounding and/or shielding might be necessary when performing electrophysiological recordings while using this device.

### Notes on safe use

1. Since direct skin contact with cold aluminium carries a potential risk of skin damage, it is essential to ensure that bed temperature does not decrease below 4°C. A thin protective layer (e.g. surgical drape) can be inserted between the animal and the metal bed to further minimize this risk, but this might result in poorer cooling efficiency. In all cases, confirming optimal temperature settings is essential before using this device on live animals.
2. While careful application and removal of surgical tape does not lead to any adverse effects in our experience, a small soft tissue pad should ideally be added between tape and animal.
3. We have not systematically assessed the maximal safe duration of cryoanesthesia using the CryoPup device. In our experience, ∼15min of cryoanesthesia result in full recovery, as observed previously by others [18]. While longer cryoanesthesia might be required for other applications, the maximal safe duration of cryoanesthesia is poorly documented and thus needs to be carefully established by experimenters.
4. For recovery, DT should be set to 21°C on the CryoPup device.
5. We recommend transferring animals to a standard recovery chamber before returning them to the maternal nest in order to fully assess their mobility.

## 8. Conclusion

We describe the design for the hardware and software of the CryoPup hypothermic anesthesia device. We provide manufacturing instructions for CryoPup and validate its use as an anesthetic device for neonatal rodents. CryoPup will enable researchers to carry out controlled hypothermic anesthesia on neonatal rodents and can seamlessly integrate into existing stereotaxic frames. This device thus provides a simple and cost-effective manner for stereotaxic surgery in neonatal rodents, facilitating the study of developing brain circuits.

## Acknowledgements

We would like to thank R. Delaqua and A. Ling from the Crick mechanical workshop for technical assistance. This work was supported by the Francis Crick Institute which receives its core funding from Cancer Research UK, the UK Medical Research Council, and the Wellcome Trust (FC001153).

## Ethics statements

Mice were bred and maintained at the specific pathogen-free mouse facility of Paris Brain Institute, with controlled temperature (21 ± 1°C) and humidity (40-70%), *ad libitum* access to standard mouse chow and water, and on a 12h:12h light/dark cycle (lights on at 8 am). Lighting cycles with a controlled rising and decreasing gradient at 8 am/8 pm respectively, simulating natural light. All procedures followed the European legislation for animal experimentation (directive 2010/63/EU).

## CRediT author statement

**BBJ:** Conceptualization, methodology, validation, formal analysis, investigation, data curation, writing – original draft, writing - review & editing, visualization; **XC-F:** conceptualization, methodology, mechanical design, electronics design, software, data curation, writing – original draft, writing - review & editing, visualization; **GK:** conceptualization, methodology, software, data curation, writing – original draft, writing - review & editing, visualization; **EdL:** Resources, validation; **NR:** Resources, validation, project administration; **AI:** writing - review & editing, project administration, supervision; **JK:** Conceptualization, methodology, resources, writing – original draft, writing - review & editing, project administration, supervision.

